# Impact of Cow Parity on the Accuracy of Near-Infrared Spectroscopy for Sustainable Milk Quality Monitoring during Milking

**DOI:** 10.64898/2026.04.13.718074

**Authors:** Patricia Iweka, Shuso Kawamura, Tomohiro Mitani

## Abstract

Accurate real-time monitoring of milk quality during milking is essential for sustainable dairy farming, yet factors such as cow parity may affect the performance of near-infrared (NIR) spectroscopic sensing systems. This study investigated how cow parity (the number of calvings) impacts the reliability and accuracy of NIR spectroscopy in assessing key milk quality indicators: fat content, lactose, and somatic cell count (SCC). Experiments were conducted at Hokkaido University with two cows in their second parity. Milk spectra were recorded across 700–1050 nm using the NIR system, while fat and lactose were measured with a MilkoScan device and SCC with a Fossomatic device. Calibration models were developed using first parity, second parity, and combined datasets through partial least squares regression. Model performance was evaluated via coefficients of determination and standard errors of prediction. Results showed comparable accuracy for milk fat and SCC across parities, whereas lactose measurements were more affected. Cross-validation between first and second parity datasets further confirmed parity-dependent variations, particularly for lactose. These findings suggest that cow parity should be considered when implementing NIR-based milk quality monitoring, supporting more precise, resource-efficient, and sustainable dairy management practices.

## 1. Introduction

Accurate determination of milk components is essential for the effective management of dairy farms as well as for industrial milk processing. The specific composition of bovine milk directly affects its suitability for various dairy products, nutritional quality, and safety standards, making it a critical factor for both producers and consumers [1–2]. Knowledge of milk constituents allows consumers to make informed choices regarding quality, while enabling processors to detect potential adulteration or inconsistencies [3]. Additionally, fluctuations in milk composition influence market value and consumer demand, which in turn affect farm income [3–4]. By assessing milk quality at the level of individual cows, farmers can implement more precise feeding strategies, monitor herd health, and optimize production efficiency, ultimately reducing resource waste and improving overall sustainability [5–8].

Near-infrared (NIR) spectroscopy has shown considerable promise as a rapid and non-destructive method for evaluating milk composition [9–13]. However, achieving accurate, real-time assessment during milking has proven challenging due to various influencing factors, including individual cow characteristics, stage of lactation, feeding practices, milking systems, seasonal variations, and cow parity [14–17]. Understanding the effect of these factors is vital for developing reliable calibration models capable of delivering high-precision measurements under practical farm conditions [4].

Enhancing real-time milk quality monitoring can contribute to more resource-efficient and environmentally responsible dairy production. By enabling precise management of feed and herd productivity, reducing milk spoilage, and supporting informed decision-making, NIR-based monitoring can minimize waste, improve profitability, and promote more sustainable dairy operations [18–19]. This study therefore investigates how cow parity influences the accuracy and reliability of NIR spectroscopic measurements of milk quality, providing insights to support sustainable and efficient dairy farm management practices.

Furthermore, implementing such precision monitoring technologies supports long-term environmental sustainability. Efficient milk quality assessment reduces overuse of feed and energy, lowers greenhouse gas emissions associated with wasted production, and promotes better animal welfare through individualized care [20–23]. By integrating these approaches into routine dairy management, farms can simultaneously achieve economic viability and ecological responsibility, demonstrating how innovative sensing technologies can contribute to a more sustainable dairy sector.

## 2. Materials and Methods

### 2.1. NIR spectroscopic sensing system

An online near-infrared (NIR) spectroscopic system was established to monitor the quality of milk from individual cows in real time during milking. The setup included an NIR spectrometer, a milk flow meter, a milk sampling unit, and a portable computer for data acquisition. The system was integrated with the milking cluster, which consisted of a claw and four teat cups, and connected to a bucket for bulk milk collection.

During milking, raw milk flowed continuously from the milking clusters into the NIR measurement chamber, holding approximately 30 mL of milk at a time, before exiting through the outlet pipe to the milk flow meter. The optical components were carefully aligned: halogen lamps A and B were positioned along the same axis as the optical fiber, while lamp C was offset by approximately 5 mm from the fiber [15].

Absorbance spectra of the milk samples were captured by the NIR spectrometer at 1 nm intervals. Measurements were recorded every 20 seconds over a wavelength range of 700–1050 nm throughout the milking session. This configuration enabled continuous, high-resolution monitoring of milk quality from individual cows.

### 2.2. Bovine and milk samples

The study was conducted using dairy cows at Hokkaido University, with measurements taken during the early lactation period to capture representative milk composition. Data collection was performed over three consecutive days, encompassing two milking sessions per day; one in the evening and the next in the morning to account for potential diurnal variations in milk quality. Milking was performed using the university’s pipeline milking system, which ensures efficient and hygienic milk collection while minimizing labor and resource use.

Milk samples were automatically gathered every 20 seconds during milking via the integrated sampling unit, allowing for high-frequency monitoring of individual cow milk quality. This approach not only provides precise and continuous data but also supports sustainable dairy management practices by reducing the need for manual sampling, lowering operational labor, and minimizing wastage of milk during quality assessment. Real-time monitoring enables targeted interventions, such as adjusting feed or managing herd health more efficiently, ultimately contributing to resource-efficient and environmentally responsible dairy production.

### 2.3. Reference (standard) analyses

To establish baseline measurements for milk quality, three key indicators were evaluated: milk fat content, lactose content, and somatic cell count (SCC). Milk fat and lactose concentrations were determined using a MilkoScan analyzer, a high-precision instrument that applies infrared spectroscopy for rapid, accurate compositional analysis. Somatic cell count, an important indicator of udder health and milk quality, was assessed using a Fossomatic device, which provides automated and reliable detection of somatic cells in milk.

These standard reference measurements were essential for calibrating the near-infrared (NIR) spectroscopic system, ensuring that the real-time monitoring data were accurate and comparable to laboratory-grade results. By using precise, automated instruments, the study minimized manual handling and measurement errors, reducing milk wastage and labor requirements. This approach aligns with sustainable dairy practices, as it supports efficient resource utilization, improved herd health management, and reduced environmental impact by allowing early detection of quality issues and targeted interventions at the individual cow level.

Overall, integrating these standard analyses with continuous NIR monitoring provides a framework for precision dairy management, enabling farms to optimize production while minimizing waste, energy consumption, and environmental footprint.

### 2.4. Calibration and validation

Three distinct datasets were obtained from two experiments corresponding to first and second parity cows. The first dataset (A) represented measurements from cows in their first parity, the second dataset (B) included cows in their second parity, and the third dataset (A+B) was a combined dataset incorporating samples from both parities.

Chemometric analyses were performed to develop and validate calibration models for each milk quality parameter—fat, lactose, and somatic cell count—and to evaluate their accuracy and precision. The analyses were conducted using The Unscrambler software (version 10.3, Camo AS, Trondheim, Norway). Both calibration and validation used all reference sample data, combining the absorbance spectra obtained from the NIR system with the corresponding laboratory reference measurements. Partial Least Squares (PLS) regression was employed to construct a calibration model for each milk parameter. The predictive accuracy of each model was assessed by comparing the NIR-predicted values to the reference measurements.

Calibration and validation were conducted under five scenarios to assess model robustness and cross-parity applicability:

1. Calibration and validation using first parity data only (A → A).
2. Calibration and validation using second parity data only (B → B).
3. Validation of second parity data using a first parity calibration model (A → B).
4. Validation of first parity data using a second parity calibration model (B → A).
5. Validation of the combined dataset using a calibration model developed from the combined dataset (A+B → A+B).

Model performance was evaluated using standard chemometric metrics, including the standard error of prediction (SEP), coefficient of determination (r^2^), bias (the mean difference between measured and predicted values), and the ratio of SEP to the standard deviation of the reference data (RPD).

By developing and validating these calibration models, the study not only ensures accurate real-time milk quality monitoring but also supports sustainable dairy management. Continuous NIR monitoring with reliable calibration models can reduce manual sampling, prevent unnecessary milk waste, optimize feed and herd management, and improve overall resource efficiency on the farm.

## 3. Results and Discussion

### 3.1. Calibration Model’s Precision and Accuracy

The validation results obtained from various test sets of milk samples for each milk quality parameter are presented in Table 1 through 5 and figure 3 to 7 below.

**Table 1.** Validation statistics for A-to-A milk quality assessment using NIR spectroscopy.

| Milk quality parameters | n | Range | $r^2$ | SEP | Bias | RPD | Regression line |
| --- | --- | --- | --- | --- | --- | --- | --- |
| Fat (%) | 72 | 1.30-7.25 | 0.99 | 0.14 | -0.00 | 11.76 | $y = 1.00x - 0.00$ |
| Lactose (%) | 72 | 3.25-4.84 | 0.87 | 0.13 | -0.00 | 2.72 | $y = 1.00x + 0.00$ |
| SCC (log SCC/mL) | 72 | 4.30-5.98 | 0.90 | 0.13 | -0.00 | 3.15 | $y = 1.00x - 0.00$ |

n, total number of validation samples; $r^2$ coefficient of determination; SEP, standard error of prediction; Bias, mean difference between measured and NIRS-predicted values; RPD, ratio of SEP to the standard deviation of reference data in the validation; regression line, regression line from predicted value (x) to reference value (y).

**Table 2.** Validation statistics for B-to-B milk quality assessment using NIR spectroscopy.

| Milk quality parameters | n | Range | $r^2$ | SEP | Bias | RPD | Regression line |
| --- | --- | --- | --- | --- | --- | --- | --- |
| Fat (%) | 78 | 1.21-9.33 | 0.99 | 0.18 | -0.00 | 11.67 | $y = 1.00x - 0.00$ |
| Lactose (%) | 78 | 3.52-4.81 | 0.78 | 0.10 | -0.00 | 2.12 | $y = 1.00x + 0.00$ |
| SCC (log SCC/mL) | 78 | 4.00-5.96 | 0.87 | 0.18 | -0.00 | 2.77 | $y = 1.00x + 0.00$ |

n, total number of validation samples; $r^2$ coefficient of determination; SEP, standard error of prediction; Bias, mean difference between measured and NIRS-predicted values; RPD, ratio of SEP to the standard deviation of reference data in the validation; regression line, regression line from predicted value (x) to reference value (y).

**Table 3.** Validation statistics for A-to-B milk quality assessment using NIR spectroscopy.

| Milk quality parameters | n | Range | r<sup>2</sup> | SEP | Bias | RPD | Regression line |
| --- | --- | --- | --- | --- | --- | --- | --- |
| Fat (%) | 78 | 1.21-9.33 | 0.98 | 0.36 | -0.28 | 5.74 | $y = 1.10x - 0.16$ |
| Lactose (%) | 78 | 3.52-4.81 | 0.43 | 0.18 | -0.06 | 1.20 | $y = 0.65x + 1.62$ |
| SCC (log SCC/mL) | 78 | 4.00-5.96 | 0.71 | 0.29 | 0.00 | 1.73 | $y = 1.32x - 1.59$ |

n, total number of validation samples; r<sup>2</sup> coefficient of determination; SEP, standard error of prediction; Bias, mean difference between measured and NIRS-predicted values; RPD, ratio of SEP to the standard deviation of reference data in the validation; regression line, regression line from predicted value (x) to reference value (y).

**Table 4.** Validation statistics for B-to-A milk quality assessment using NIR spectroscopy.

| Milk quality parameters | n | Range | r<sup>2</sup> | SEP | Bias | RPD | Regression line |
| --- | --- | --- | --- | --- | --- | --- | --- |
| Fat (%) | 72 | 1.30-7.25 | 0.95 | 0.43 | 0.04 | 3.87 | $y = 0.87x + 0.51$ |
| Lactose (%) | 72 | 3.25-4.84 | 0.41 | 0.28 | 0.05 | 1.26 | $y = 1.46x - 2.09$ |
| SCC (log SCC/mL) | 72 | 4.30-5.98 | 0.52 | 0.31 | 0.00 | 1.37 | $y = 0.76x + 1.21$ |

n, total number of validation samples; r<sup>2</sup> coefficient of determination; SEP, standard error of prediction; Bias, mean difference between measured and NIRS-predicted values; RPD, ratio of SEP to the standard deviation of reference data in the validation; regression line, regression line from predicted value (x) to reference value (y).

**Table 5.** Validation statistics for A*+*B milk quality assessment using NIR spectroscopy.

| Milk quality parameters | n | Range | r<sup>2</sup> | SEP | Bias | RPD | Regression line |
| --- | --- | --- | --- | --- | --- | --- | --- |
| Fat (%) | 150 | 1.21-9.33 | 0.98 | 0.25 | -0.00 | 7.69 | $y = 1.00x - 0.00$ |
| Lactose (%) | 150 | 3.25-4.84 | 0.59 | 0.19 | -0.00 | 1.55 | $y = 1.00x + 0.00$ |
| SCC (log SCC/mL) | 150 | 4.00-50.98 | 0.79 | 0.21 | -0.00 | 2.16 | $y = 1.00x + 0.00$ |

n, total number of validation samples; r<sup>2</sup> coefficient of determination; SEP, standard error of prediction; Bias, mean difference between measured and NIRS-predicted values; RPD, ratio of SEP to the standard deviation of reference data in the validation; regression line, regression line from predicted value (x) to reference value (y).

**Figure 1.**
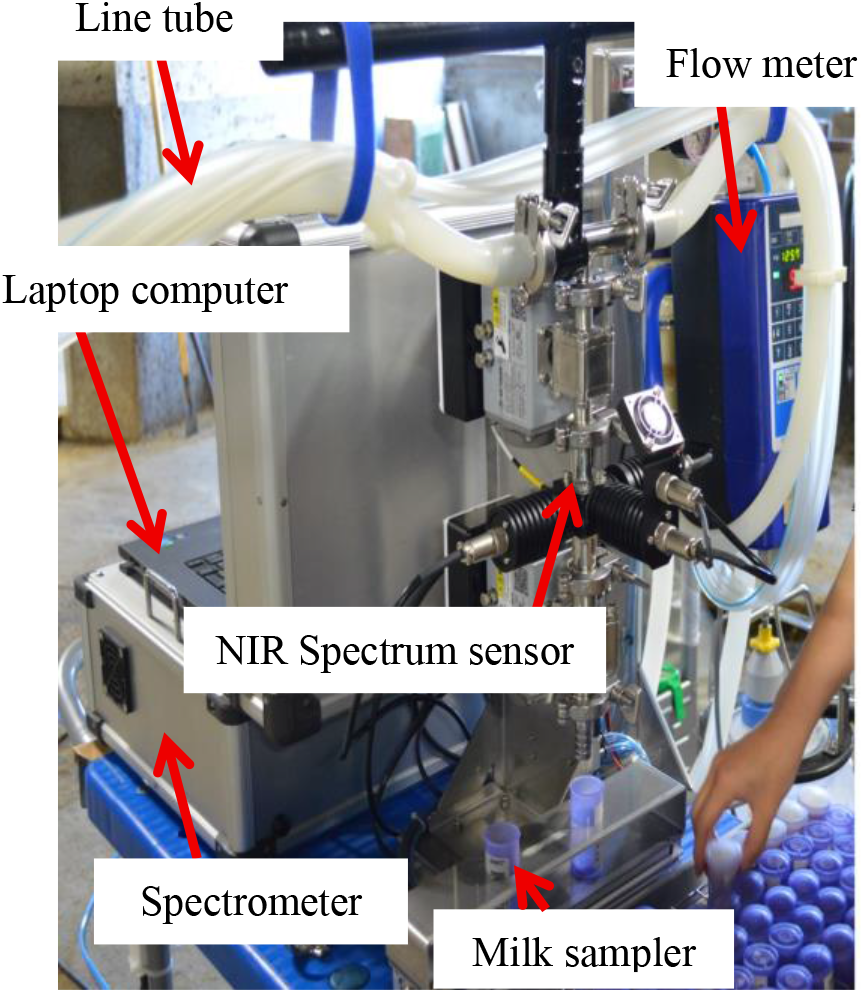
Online real-time NIR sensing system

**Figure 2.**
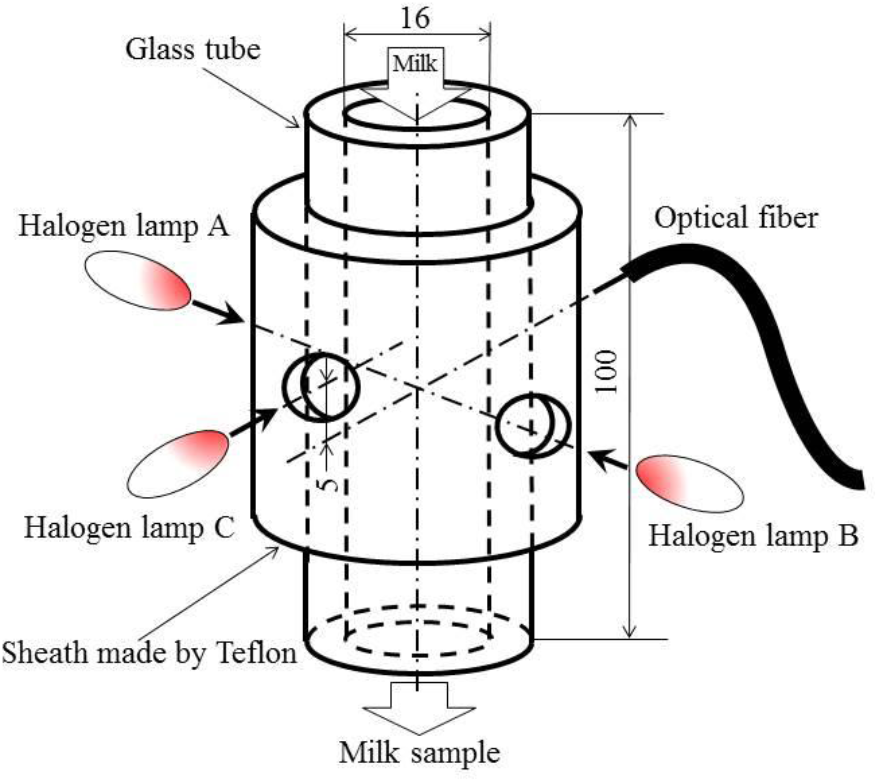
Diagram of the NIR spectrum sensor’s milk chamber optical system

**Figure 3.**
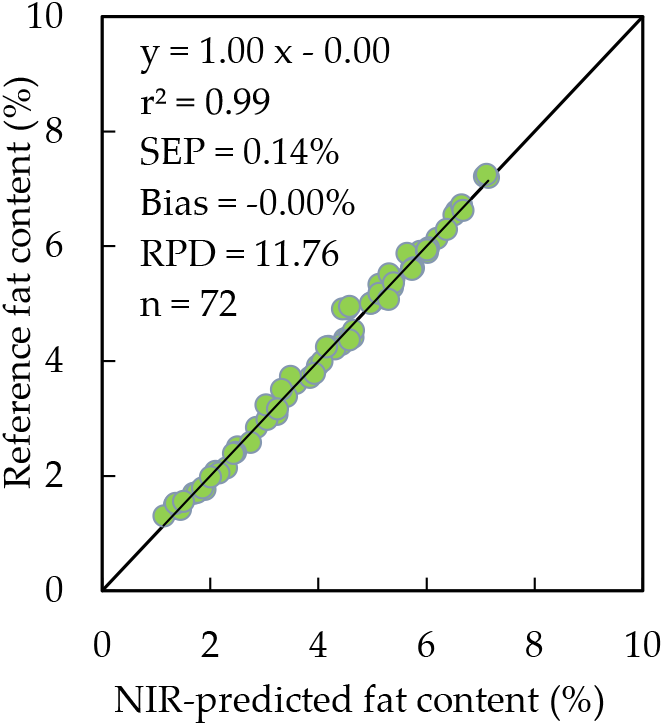
Relationship between measured fat content and NIR-estimated fat content (1^st^ parity, A to A)

**Figure 4.**
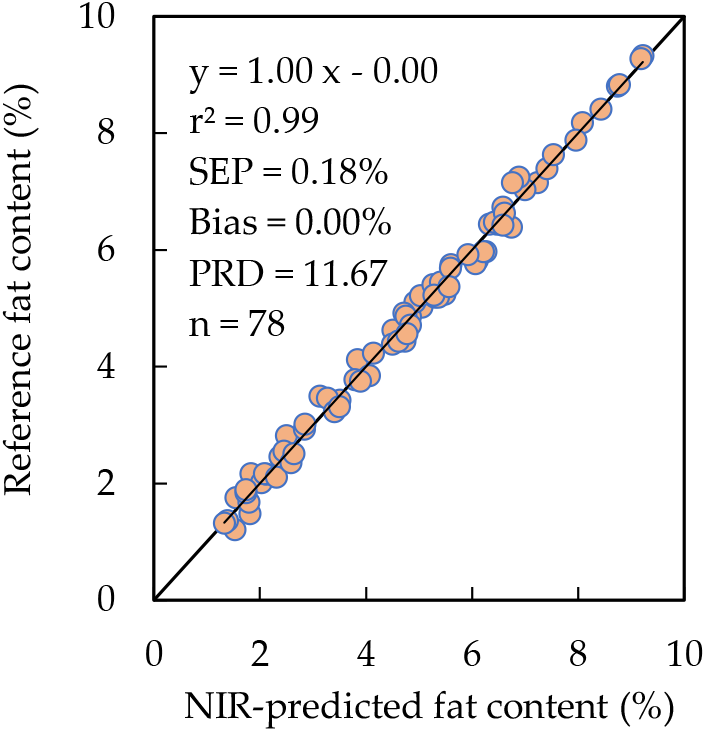
Relationship between measured fat content and NIR-estimated fat content (2^nd^ parity, B to B)

**Figure 5.**
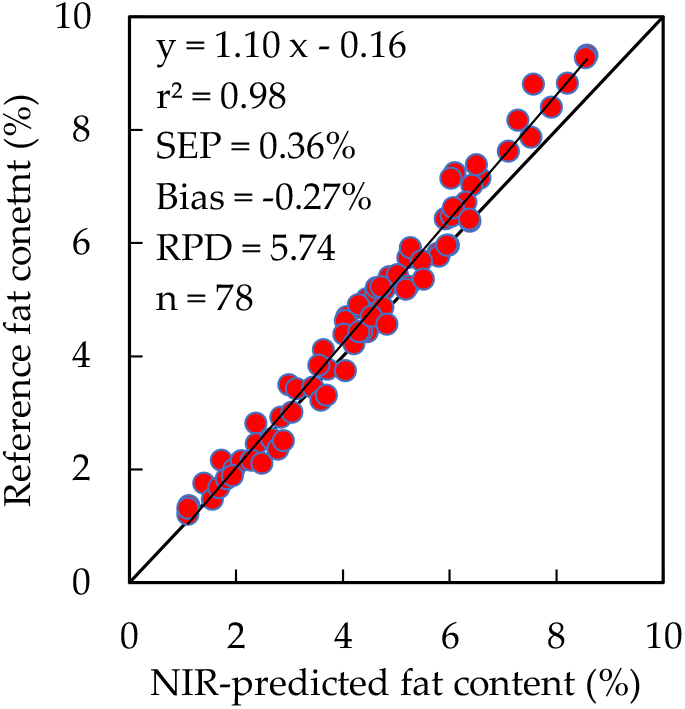
Relationship between measured fat content and NIR-estimated fat content(1^st^ vs 2^nd^parity, A to B)

**Figure 6.**
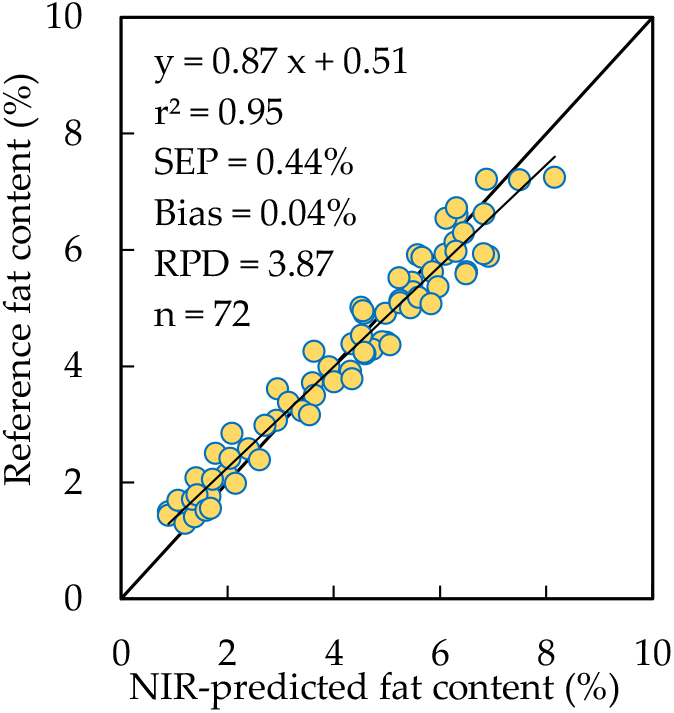
Relationship between measured fat content and NIR-estimated fat content(2^st^ vs 1^nd^parity, A to B)

**Figure 7.**
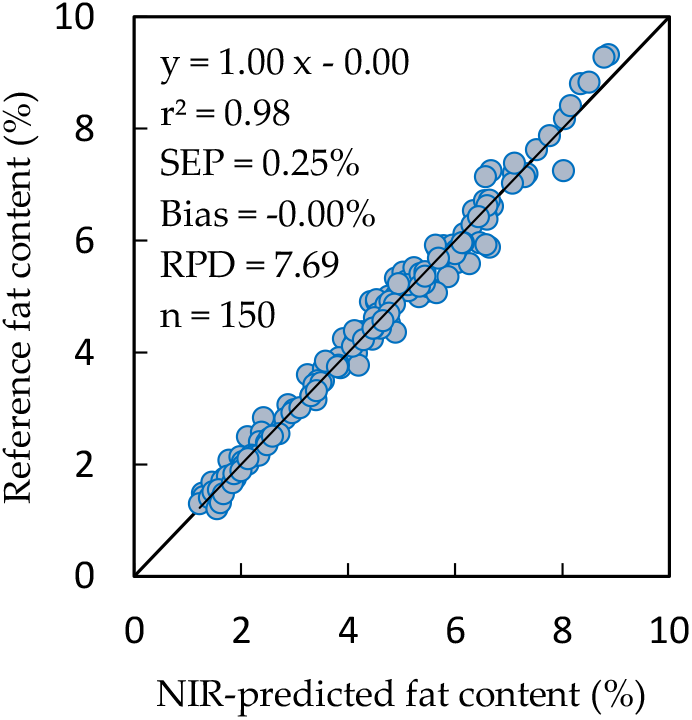
Relationship between measured fat content and NIR-estimated fat content (1^st &^ 2^nd^ parity, A + B)

Calibration models developed using first-parity cow samples (A → A) exhibited high predictive accuracy for milk fat, lactose, and somatic cell count (SCC). The standard error of prediction (SEP) for fat and lactose was below 1%, while SEP for SCC remained under 1 log SCC/mL. Minimal bias values indicated negligible systematic error, and high coefficients of determination (r^2^) reflected reliable model performance. These results confirm that NIR spectroscopy can effectively monitor milk quality within a single parity group, providing rapid and accurate compositional data.

Models constructed from second-parity samples (B → B) similarly showed strong predictive performance across all three parameters, with low SEP and high r^2^ values, confirming the robustness of the NIR calibration approach. Comparison of single-parity models indicated a slight increase in SEP for second-parity predictions, suggesting minor compositional variability within this group, though bias remained minimal.

Cross-parity validation revealed notable declines in model performance. First-parity calibration models applied to second-parity milk (A → B) exhibited decreased r^2^ and increased SEP, particularly for lactose, while the reverse scenario (B → A) showed similar reductions in accuracy and increased bias. These findings suggest that parity-related variations in milk composition, rather than random errors, are the primary contributors to reduced predictive reliability across parities. Such variations are driven by physiological and metabolic differences: medium- and long-chain fatty acids generally increase with parity due to higher circulating triglycerides, while first-parity cows allocate more nutrients to growth rather than milk fat synthesis. Additionally, parity influences milk protein, casein, and corticosteroid levels, all of which modify chemical properties and the NIR absorbance spectra.

Calibration models developed from a combined dataset of first- and second-parity cows (A + B) improved predictive performance across both groups, demonstrating that including multiple parity groups enhances model robustness. Although SEP increased slightly and r^2^ decreased compared to single-parity models, the combined dataset reduced the large errors observed in cross-parity validations. These results emphasize the importance of constructing inclusive calibration models that account for physiological and compositional variability among cows, particularly when aiming for broad applicability in commercial dairy herds.

The findings have multiple practical benefits. Accurate, real-time monitoring of milk quality supports precision feeding, allowing farmers to allocate nutrients efficiently according to individual cow needs, reducing overfeeding and minimizing waste. Early detection of deviations in milk composition or SCC enables timely intervention, preventing milk spoilage, reducing antibiotic use, and improving animal welfare. Optimized herd management can enhance milk yield and composition consistency, ultimately lowering the environmental footprint by minimizing greenhouse gas emissions and resource consumption per liter of milk produced.

Moreover, incorporating parity-specific information into NIR calibration models contributes to data-driven, sustainable dairy management. By capturing variations across parity groups, farms can implement more targeted strategies for feeding, health monitoring, and lactation management. This not only enhances predictive accuracy but also promotes economic sustainability through improved productivity and reduced losses, while advancing ecological sustainability by conserving feed, water, and energy resources.

The study demonstrates that parity significantly influences NIR-based milk quality predictions, particularly for lactose, and highlights the critical role of tailored calibration models in achieving precision dairy farming. These insights provide a practical framework for developing sustainable, efficient, and environmentally responsible milk production systems, where real-time compositional monitoring drives optimized resource use, reduced waste, and improved herd health outcomes.

## 4. Conclusions

This study demonstrates that cow parity significantly influences the accuracy of near-infrared (NIR) spectroscopic models used for real-time monitoring of milk quality parameters, particularly lactose. Calibration models developed from single-parity datasets (first or second parity) performed poorly when applied to milk from cows of a different parity, highlighting the necessity of considering parity-related variations in milk composition. Including data from multiple parities in calibration models markedly improves predictive accuracy, precision, and robustness across different groups of cows.

From a sustainability standpoint, these findings have substantial implications for modern dairy management. Accurate, real-time milk quality monitoring allows farmers to make data-driven decisions regarding feed allocation, herd management, and milking practices, minimizing unnecessary resource use. By identifying milk quality deviations promptly, farmers can reduce milk waste, optimize energy consumption in milk processing, and prevent overfeeding or underfeeding of cows, which in turn lowers greenhouse gas emissions and environmental footprint. Furthermore, precise monitoring supports animal health and welfare, as deviations in milk composition can indicate mastitis or metabolic stress, allowing early intervention and reducing the need for intensive treatments.

Incorporating parity-specific calibration models into NIR systems contributes to precision dairy farming, enabling farms to enhance both economic efficiency and ecological responsibility. By optimizing resource utilization and minimizing waste, this approach promotes sustainable milk production that balances profitability, environmental stewardship, and animal welfare. Overall, this study underscores the critical role of integrating parity-based considerations into milk quality monitoring systems and provides a practical framework for advancing sustainable, data-driven dairy farming practices.

## Author Contributions

Conceptualization, S.K. and P.I.; methodology, S.K., P.I., and T.M.; software, S.K. and P.I.; validation, P.I. and S.K.; formal analysis, P.I., S.K., and T.M.; investigation, P.I., S.K., and T.M.; data curation, P.I. and S.K.; writing-original draft preparation, P.I.; writing review and editing, S.K., T.M., and P.I. All authors have reviewed and approved the final version of the manuscript for publication.

## Institutional Review Board Statement

Not applicable.

## Informed Consent Statement

Not applicable.

## Data Availability Statement

This paper presents a summary of all the new data acquired during the study.

